# Establishing the characteristics of an effective pharmacogenetic test for clozapine induced agranulocytosis

**DOI:** 10.1101/010900

**Authors:** Moira Verbelen, David A Collier, Dan Cohen, James H MacCabe, Cathryn M Lewis

## Abstract

Clozapine is the only evidence-based therapy for treatment resistant schizophrenia, but it induces agranulocytosis, a rare but potentially fatal haematological adverse reaction, in less than 1% of users. To improve safety, the drug is subject to mandatory haematological monitoring throughout the course of treatment, which is burdensome for the patient and one of the main reasons clozapine is underused. Therefore, a pharmacogenetic test is clinically useful if it identifies a group of patients for whom the agranulocytosis risk is low enough to alleviate monitoring requirements. Assuming a genotypic marker stratifies patients into a high risk and a low risk group, we explore the relationship between test sensitivity, group size and agranulocytosis risk. High sensitivity minimizes the agranulocytosis risk in the low risk group and is essential for clinical utility, in particular in combination with a small high risk group.

## Introduction

Although there are 22 FDA approved antipsychotics used to treat schizophrenia, around 30% of schizophrenia patients do not respond to drugs other than clozapine.^1^ Clozapine has superior efficacy for positive symptoms in these treatment resistant patients^2^ and may improve negative symptoms.^3^ Furthermore, clozapine reduces suicidal behaviour especially when compared with first generation antipsychotics and overall mortality at population level.^4-8^

Despite its proven efficacy, the clinical use of clozapine is limited by the risk of agranulocytosis, a rare but potentially fatal adverse drug reaction, characterised by the acute loss of neutrophils in circulating blood. Agranulocytosis is defined as an absolute neutrophil count (ANC) of less than 500 cells/mm³ blood. Shortly after clozapine was introduced in Europe in the 1970s, it was withdrawn from the market when 17 cases of agranulocytosis were reported in Finland, of which 8 were fatal.^9^ In 1990, clozapine was reintroduced after its superiority over chlorpromazine for the treatment of refractory schizophrenia was shown.^10^ However, its use was restricted in most Western countries to treatment of refractory patients, i.e. patients who have not improved on at least two different antipsychotics.^2, 11-13^

In order to prevent agranulocytosis by detecting a fall in ANC, patients treated with clozapine are subject to compulsory haematological monitoring. In Europe, the full white blood cell count and ANC are monitored weekly for the first 18 weeks of treatment and every 4 weeks thereafter for the duration of the treatment.^14^ If at any time during treatment the white blood cell count falls below 3000cells/mm³ or the ANC below 1500 cells/mm³, clozapine should be discontinued immediately and these patients should not be treated with clozapine again except in a controlled setting.^15-17^ Although the obligatory monitoring has the benefit of regular contact with a health care professional, it is an invasive procedure and can be a burden for the patient. Moreover, some patients decline to take clozapine because of the monitoring requirement.^18^

As agranulocytosis can develop within 2 to 5 days even weekly monitoring cannot guarantee timely detection in all cases from occurring.^19^ The incidence of agranulocytosis induced by clozapine varies between 0.38% and 0.8%, with approximately 80% of cases occurring within the first 18 weeks.^20-23^ The incidence of agranulocytosis decreases from 0.7% in the first year, to 0.07% or lower in the second year of treatment.^24, 25^ Few cases occur later in the course of treatment but the risk does not fully disappear. In 2 to 4% of patients agranulocytosis is fatal, which corresponds to an overall mortality rate of about 1 to 3 in 10000 patients on clozapine.^26^ However, most patients recover completely from agranulocytosis with no haematological consequences.^23, 24, 27^

In spite of its therapeutic advantages with respect to its efficacy in treatment resistant schizophrenia, clozapine is underused, mainly due to the risk of severe adverse events, primarily agranulocytosis, and the mandatory haematological monitoring.^28^ Around 30% of schizophrenia patients meet the indications for clozapine treatment, but the market share of clozapine, which is now a generic drug, was less than 5% in 2010 in the US.^2^

A pharmacogenetic test for clozapine induced agranulocytosis could greatly improve the burden of haematological monitoring if the monitoring requirements could be made less onerous, or be time limited, for the majority of patients with a low genetic risk for agranulocytosis. Not only would this make clozapine treatment more acceptable for the patient, it would also save considerable health care resources. On the other hand, the patients who are at higher risk of developing agranulocytosis could be monitored more frequently or, if the risk is very high, not exposed to clozapine at all.

Pharmacogenetic research of clozapine induced agranulocytosis has focused on candidate genes in case-control studies. Several associations with human leukocyte antigen (HLA) alleles have been reported, as well as associations with the tumour necrosis factor and N-ribosyldihydronicotin-amide quinone oxido-reductase 2 (NQO2) genes.^29, 30^ However, few of these findings have been replicated and the majority of these pharmacogenetic studies suffered from typical candidate gene study issues, namely small sample sizes and inadequate correction for multiple testing. The most promising finding was that the HLA-DQB1 6672G>C polymorphism was associated with clozapine induced agranulocytosis, with an odds ratio of 16.9.^31^ A pharmacogenetic test based on this polymorphism has been marketed, but due to low sensitivity (21.5%) it failed to be a commercial or clinical success.^29, 32^ In the first genome wide association study, amino acid changes in HLA-DQB1 (126Q) and HLA-B (158T) were associated with clozapine induced agranulocytosis with more modest odds ratios of 0.19 and 3.11 respectively.^33^

Here we investigate the required properties of a clinically useful pharmacogenetic test that could stratify clozapine users with regards to their agranulocytosis risk as described above.

## Methods

We assume that the genetic test divides patients into two groups with different levels of agranulocytosis risk, and that the low risk (LR) group contains a higher proportion of patients than the high risk (HR) group. When comparing the outcome of a pharmacogenetic agranulocytosis test with the actual agranulocytosis status, the following scenarios can occur (Table 1):

- True positive: A patient who does develop agranulocytosis is correctly identified as HR. This scenario has probability *a*.
- False positive: A patient who does not develop agranulocytosis is wrongly identified as HR. This scenario has probability *b*.
- False negative: A patient who does develop agranulocytosis is wrongly identified as LR. This scenario has probability *c*.
- True negative: A patient who does not develop agranulocytosis is correctly identified as LR. This scenario has probability *d*.

**Table 1.**
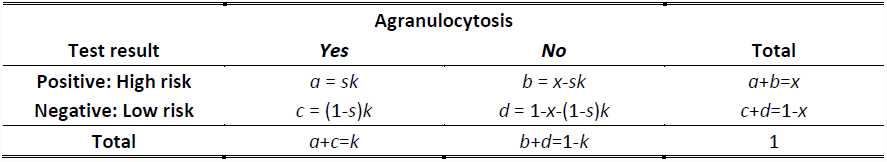
Classification table comparing test outcome with true agranulocytosis status. Cell probabilities are expressed in terms of sensitivity (*s*) and size of the HR group (*x*).

If the incidence of clozapine induced agranulocytosis is a known parameter *k*, the agranulocytosis risk regardless of test outcome (*a*+*c*) equals *k* and the probability of not getting agranulocytosis (*b*+*d*) is 1-*k*.

Assuming that *k* is known, two of the following parameters need to be fixed in order to calculate the probabilities in each cell of Table 1:

- The proportion of patients in the HR group (*x*) or the proportion of patients in the LR group (1-*x*).
- The sensitivity of the test, which is the proportion of correctly classified agranulocytosis cases. In Table 1,

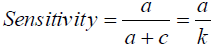
- The specificity of the test, which is the proportion of correctly classified agranulocytosis free patients. In Table 1,

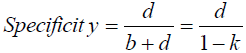

The proportion of patients in each risk group (*x*, 1-*x*) is relevant to this study, as we want to justify a more lenient monitoring schedule for the LR group. The larger the LR group is the more patients on clozapine will benefit from monitoring regime changes. Assuming a single locus test, the size of the two risk groups depends on the allele frequency of the test marker. Hence, we use the proportion of patients in the HR group (x) and test sensitivity (*s*) to study the cell probabilities in Table 1.

Of primary interest is the agranulocytosis risk in the LR group, as these are the patients for whom the haematological monitoring rules could be relaxed, and this outcome corresponds to the complement of the negative predictive value (NPV), being the proportion of test negative or LR patients who do not develop agranulocytosis. Therefore, we investigate the relationship between agranulocytosis risk in the LR group, test sensitivity and the size of the HR group. The agranulocytosis risk in the LR group is given by

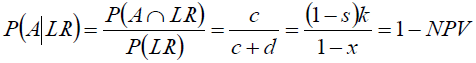

where *A* stands for developing agranulocytosis. Thus, the agranulocytosis risk in the LR group decreases as test sensitivity increases or as the HR group becomes smaller, which corresponds to the LR group getting larger.

As a secondary outcome, we study the agranulocytosis risk in the HR group, which corresponds to the positive predictive value (PPV) or the proportion of test positive patients who are true agranulocytosis cases, and is given by

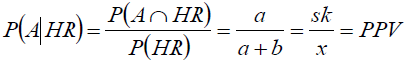

In the HR group, the agranulocytosis risk increases when sensitivity rises or the proportion of patients in the HR group decreases.

By definition the agranulocytosis risk in the HR group must be larger than in the LR group, so

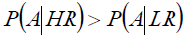

or expressed in terms of sensitivity and size of the HR group

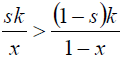

which reduces to *s* > *x*. In other words, sensitivity must be larger than the proportion of patients in the HR group.

We explore the relationship between different parameters, focusing on sensitivity (*s*), and the proportion of patients assigned to the HR group (*x*), in the context of a genetic test to predict the risk of clozapine induced agranulocytosis. Furthermore, we develop guidelines for a test that divides the population into a LR and HR group, assuming the total risk of developing clozapine induced agranulocytosis(*k*) is 0.8%.^11^ We also assess how the pharmacogenetic test based on the HLA-DQB1 6672G>C polymorphism performs under this framework.^31^

## Results

The key parameters for a clinically effective test, i.e. a test that minimizes the agranulocytosis risk in the LR group, are high sensitivity and to a lesser extent a small proportion of patients assigned to the HR group. Figure 1 shows that to obtain a low agranulocytosis risk in the LR group (solid lines), with a concomitant high risk in the HR group (dotted lines), a test should be highly sensitive. Also, for a given sensitivity, a smaller HR group corresponds to lower agranulocytosis risk in the LR group and a higher agranulocytosis risk in the HR group (Figure 1 and Table2). The lower the sensitivity of a test is, the smaller the difference between the agranulocytosis risks in both groups. When the sensitivity is equal to the size of the HR group, the risk in the two groups are the same and equal to the overall agranulocytosis risk of 0.8%. A test with sensitivity close to the proportion of patients in the HR group would thus be irrelevant.

**Table 2.** Hypothetical tests with 80% sensitivity for different proportions of patients in the HR group, showing agranulocytosis risks in the LR group (1-NPV), the HR group (PPV) and specificity of the test.

| **Size of HR group** | **P(A \| LR)**<br>**(1-NPV)** | **P(A \| HR)**<br>**(PPV)** | **Specificity** |
| --- | --- | --- | --- |
| 0.010 | 0.0016 | 0.640 | 0.996 |
| 0.050 | 0.0017 | 0.128 | 0.956 |
| 0.100 | 0.0018 | 0.064 | 0.906 |
| 0.200 | 0.0020 | 0.032 | 0.805 |
| 0.300 | 0.0023 | 0.021 | 0.704 |
| 0.400 | 0.0027 | 0.016 | 0.603 |
| 0.500 | 0.0032 | 0.013 | 0.502 |

**Figure 1.**
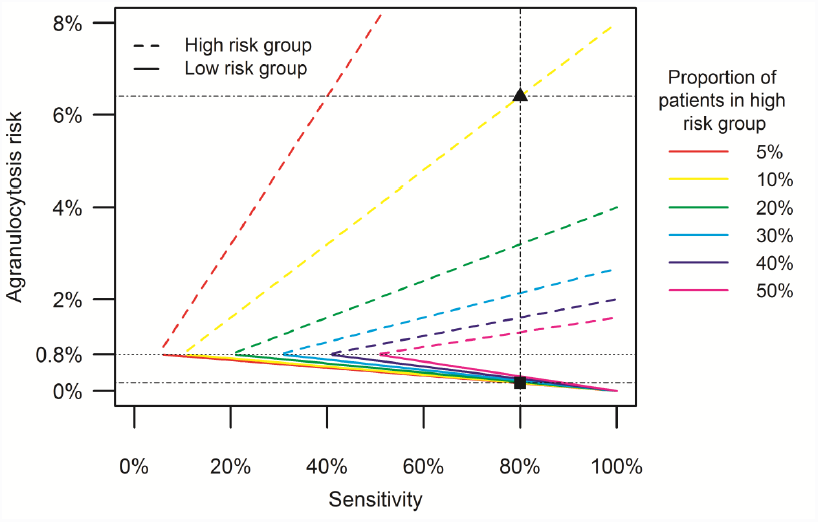
Agranulocytosis risk in LR and HR groups by sensitivity, for different HR group sizes between 5% and 50%. The ■ and ▲ indicate the agranulocytosis risk in the LR and HR groups, respectively, for a test with 80% sensitivity and 10% of patients in the HR group.

Exploration of the relationship between agranulocytosis risk and the proportion of patients in the HR group confirms that a smaller HR group gives rise to a lower agranulocytosis risk in the LR group (Figure 2). The risk in the HR group increases steeply when the size of that group is close to zero. For a given size of the HR group, high sensitivity leads to a low agranulocytosis risk in the LR group and a high risk in the HR group (Figure 2 and Table 3). As in Figure 1, the risk curves in Figure 2 meet at 0.8% agranulocytosis risk when the proportion of patients in the HR group is equal to the sensitivity (except for 100% sensitivity where the agranulocytosis risk in the LR group is zero as all agranulocytosis cases are detected by the test).

**Table 3.**
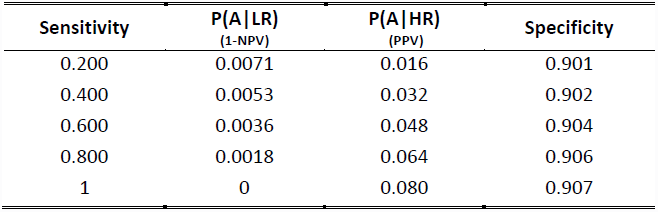
Hypothetical tests with 10% of patients in the HR group for different sensitivity values, showing agranulocytosis risks the LR group (1-NPV), in the HR group (PPV) and specificity of the test.

**Figure 2.**
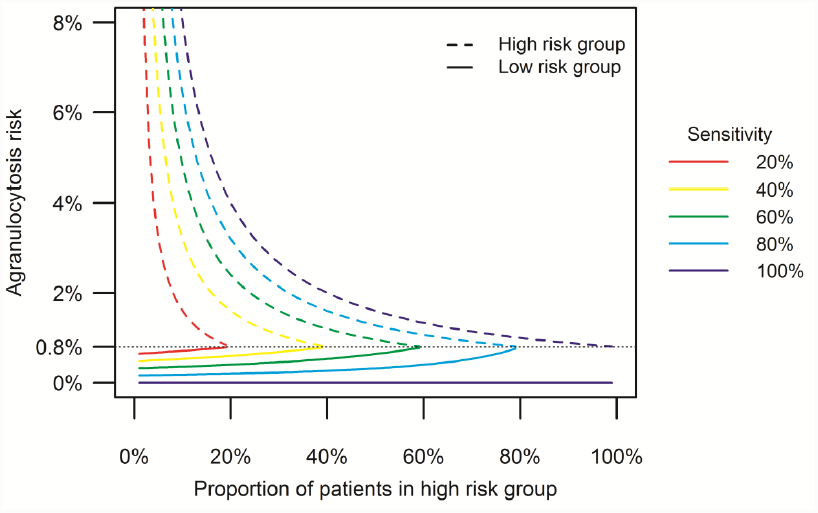
Agranulocytosis risk in LR and HR groups by proportion of patients that classified as HR, for different sensitivity values between 20% and 100%.

A simultaneous assessment of sensitivity and HR group size shows that test sensitivity controls the reduction in agranulocytosis risk seen in the LR group, with high sensitivity leading to low agranulocytosis risk (Figure 3a). High sensitivity implies that most agranulocytosis cases are identified by the genetic test and classified as HR. By consequence nearly all patients in the LR group do not develop agranulocytosis, and hence the risk in that group is low. For example, if we need the agranulocytosis risk in the LR group to be half the population risk, (i.e. ≤ 0.4%), the test sensitivity must be at least 50.2%. To achieve a stratification where the LR group is at one fifth of the average agranulocytosis risk, sensitivity greater than 80.1% is required. A small HR group contributes to a low agranulocytosis risk in the LR group by preventing true LR patients from being wrongly classified as HR. A large number of true LR patients maximizes the denominator of the agranulocytosis risk in the LR group, and thus minimizes the risk itself.

**Figure 3.**
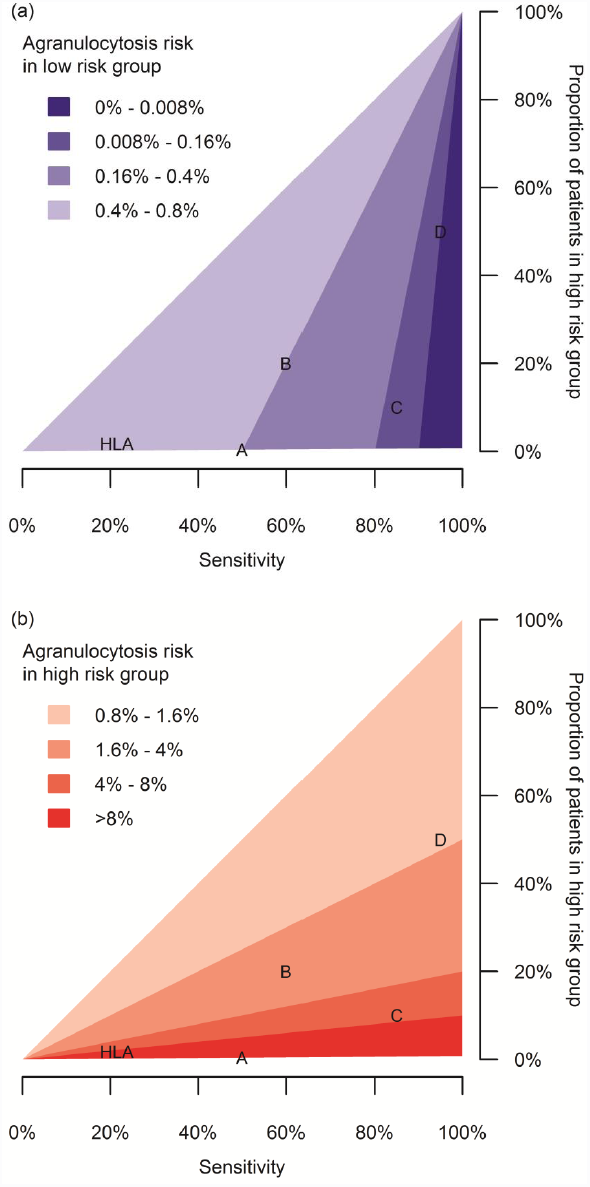
Agranulocytosis risk in (a) LR group and (b) HR group by sensitivity and proportion of patients in the HR group. In both panels, darker colours represent the desired outcomes of low risk in LR group and high risk in the HR group. The letters indicate the position of hypothetical tests A, B, C and D; HLA indicates the position of the HLA-DQB1 6672G>C based test.

The agranulocytosis risk in the HR group depends largely on the size of the HR group (Figure 3b). In a smaller HR group, the ratio of true HR patients versus patients incorrectly classified as HR is larger, and so the agranulocytosis risk in the HR group is larger. High sensitivity increases the number of true HR patients and in that way leads to a high agranulocytosis risk in this group, but even in the ideal scenario of maximum sensitivity the proportion of true HR patients is limited to 0.8%.

High sensitivity and a small HR group lead to an effective test with the small group of HR patients requiring more frequent monitoring, while the majority of patients are assigned to the LR group, which has substantially reduced agranulocytosis risk and could therefore be monitored less frequently.

To comment on the clinical utility of a pharmacogenetic test, an agranulocytosis risk that is acceptable without monitoring must be determined. We propose that an agranulocytosis risk of 0.13% is acceptable, because this corresponds to the risk conferred by the antipsychotic chlorpromazine which does not have mandatory monitoring in the UK.^34, 35^ To achieve this, the sensitivity of the test must be at least 83.9%. We examine four hypothetical pharmacogenetic tests and how the outcomes affect haematological monitoring of patients (Table 4).

- Test A has 50% sensitivity, which is the maximum sensitivity that can be achieved with such a small HR group (0.4%). All HR patients will develop agranulocytosis, so test A is useful to prevent agranulocytosis in the small group of HR patients by not exposing them to clozapine, but the remaining 99.6% of patients have an agranulocytosis risk that is too high to reduce monitoring. This test is thus only clinically relevant for a small proportion of patients.
- Test B has a higher sensitivity and a larger HR group than test A, and results in the same agranulocytosis risk for the LR group. However, the risk in the HR group is much lower than for test A. Considering there is no alternative to clozapine, it is less clear whether not to treat the HR patients with clozapine. As there is no definite impact on the treatment of either risk group, test B is not clinically useful.
- Test C has a relatively high sensitivity of 85% and classifies 10% of patients as HR. The agranulocytosis risk in the LR group is 0.13%, a risk we argue is low enough to stop or reduce haematological monitoring. Nine out of 10 patients would benefit from test C in this way.
- Test D results in an even lower agranulocytosis risk for the LR group, but fewer patients would benefit from changes in the monitoring schedule as the LR group contains only 50% of patients. Although the sensitivity of test D is higher than of test C, the smaller size of the HR group in test C means that this test would have most clinical impact.

**Table 4.** Four hypothetical pharmacogenetic tests for clozapine induced agranulocytosis and their clinical impact.

| Test | Sensitivity | Size of HR group | P(A\|LR)<br>(1-NPV) | P(A\|HR)<br>(PPV) | Specificity | Clinical impact |
| --- | --- | --- | --- | --- | --- | --- |
| A | 0.500 | 0.004 | 0.004 | 1 | 1 | No change in monitoring, but withhold clozapine from HR group |
| B | 0.600 | 0.200 | 0.004 | 0.024 | 0.803 | No change in monitoring |
| C | 0.850 | 0.100 | 0.0013 | 0.068 | 0.906 | Stop monitoring in LR group |
| D | 0.950 | 0.500 | 0.0008 | 0.015 | 0.504 | Stop monitoring in LR group |

The pharmacogenetic agranulocytosis test using the HLA-DQB1 6672G>C polymorphism has a sensitivity of 21.5% and specificity of 98.4%.^31^ Based on those values and assuming an overall agranulocytosis risk of 0.8%, the test classifies 1.76% of patients as HR, with a high agranulocytosis risk of 9.66%. Conversely, the agranulocytosis risk in the LR group is 0.64%. This is a relative risk of 0.8 or a 20% reduction in risk compared with the agranulocytosis risk without genetic stratification, and exceeds our maximum acceptable agranulocytosis risk of 0.13%. Hence, the HLA-DQB1 6672G>C based genetic test has limited use in stratifying patients in order to reduce haematological monitoring requirements for a subset of patients.

## Discussion

We have established a framework for assessing the utility of a genetic test for clozapine-induced agranulocytosis and explored the characteristics of tests that would reduce agranulocytosis risk to a level which does not require regular haematological monitoring. In particular, we show that high sensitivity is essential and that a small proportion of patients classified as HR further decreases the agranulocytosis risk in the LR group.

High sensitivity is a self-evident characteristic of a clinically useful test, but the finding that a small HR group is favourable might seem counterintuitive. One could reason that to be sure the LR group contains no agranulocytosis cases, the HR group should include all patients who are at the slightest risk of developing agranulocytosis, and that the HR group should thus be large. However, since the number of true agranulocytosis cases is very low, a large HR group would mainly contain false positive patients. Instead of a large HR group, a useful test relies on high sensitivity to correctly classify the agranulocytosis cases as HR. A small HR group implies few false positives and many true negatives, which in turn minimizes the agranulocytosis risk in the LR group.

Instead of a single genetic locus, a pharmacogenetic test could also be based on polygenic risk scores, built from combining risk conferred by many genetic loci to identify patients at high risk. In that case, the threshold defining LR and HR groups can be varied, and the effectiveness of a test is typically measured by the area under a receiver operating characteristic curve (AUC).^36^ When the threshold is moved to increase test sensitivity, the size of the HR group size will increase as well. It is not straightforward to predict the resulting change – increase or decrease - in agranulocytosis risks in the LR and HR groups, because these depend on the distribution of the polygenic risk scores. Once an appropriate polygenic score threshold has been fixed, a test can be translated easily to the framework developed here.

Some pharmacogenetic tests for other drugs show high sensitivity and are in clinical use. Genetic associations of HLA-alleles that predict adverse drug reactions caused by abacavir and carbamazepine have sensitivity and specificity values close to 100% (Table 5). About 4% of patients treated with abacavir, a HIV-1 reverse transcriptase inhibitor, develop a potentially fatal hypersensitivity syndrome.^37, 38^ The clinical usefulness of HLA-B*5701 testing prior to abacavir initiation was confirmed in a clinical trial, which found that prospective HLA-B*5701 screening reduced the incidence of hypersensitivity syndrome compared with no screening.^39, 40^ All hypersensitivity cases had the HLA-B*5701 allele but nearly half of HLA-B-5701 carriers did not develop this adverse drug reaction. In the UK and the US, screening for HLA-B*5701 is recommended before starting abacavir treatment.^41, 42^

**Table 5.** Characteristics of pharmacogenetic tests that predict adverse drug reactions.

| **Drug** | **Gene** | **Test** | **Performance characteristics** | **Reference** |
| --- | --- | --- | --- | --- |
| Abacavir | HLA-B*5701 | The HLA-B*5701 allele is associated with hypersensitivity reaction. Carriers should avoid abacavir. | Sensitivity: 100%<br>Specificity: 96.9%<br>PPV: 47.9%<br>NPV: 100% | 39 |
| Carbamazepine | HLA-B*1502 | The HLA-B*1502 allele is associated with SJS-TEN in Han Chinese. Carriers should avoid carbamazepine. | Sensitivity: 98.3%<br>Specificity: 97%<br>PPV: 7.7%<br>NPV: 100% | 44 |
| Purine analogues | TPMT | Impute 2 SNPs to identify patients with zero wildtype alleles, as they are at high risk of myelotoxicity. | Sensitivity: 100%<br>Specificity: 99.0%<br>PPV: 80.6%<br>NPV: 100% | 48 |

The antiepileptic drug carbamazepine can cause the severe life-threatening skin reactions Stevens-Johnson Syndrome (SJS) and toxic epidermal necrolysis (TEN), with a mortality rate of 30-40%.^43, 44^ In Asian populations, the HLA-B*1502 allele confers risk of SJS-TEN with high sensitivity and specificity.^43, 45, 46^ This marker picks up nearly all SJS-TEN cases, but many HLA-B*1502 positive patients do not develop SJS-TEN, hence the low PPV. HLA-B*1502 testing is required by the FDA for Asian patients who are prescribed carbamazepine.^44, 47^

Recently, the imputation of thiopurine methyltransferase (TPMT) alleles was described as a method to predict TPMT enzyme activity with high sensitivity and specificity.^48^ Purine analogues such as azathioprine, 6-mercaptopurine and thioguanine are metabolized by TPMT, an enzyme for which there is considerable genetic variability in activity.^49^ The FDA recommends that patients are genotyped or phenotyped for TPMT activity before starting purine analogue treatment in order to adjust the starting dose.^50, 51^ Moreover, patients with no functional allele and thus zero TPMT activity could be offered an alternative drug, as they are at very high risk of developing severe leukopenia and myelosuppression.^52^ TPMT activity can be assessed by an enzyme activity test or by imputing two SNPs that represent 3 defective TPMT alleles. SNP imputation after genetic testing identified null-TPMT activity individuals with very high sensitivity and specificity. All zero TPMT activity individuals are detected by the test, but around 1 in 5 test positives are false positive.^48^

The sensitivity, specificity and NPV of these genetic tests are all close to 100%. On the other hand, the PPV values vary between 7.7% and 80.6%. These low PPVs are acceptable, since there are alternative treatments available for abacavir, carbamazepine and purine analogues, and patients who test positive can be treated with a different drug. In contrast, clozapine is reserved for treatment of refractory schizophrenia and no alternative drug is available. A high PPV would ensure that few patients are unnecessarily excluded from treatment, but a high NPV and consequently low agranulocytosis risk is also important to justify a reduced monitoring schedule for patients in the LR group.

No genetic test for clozapine induced agranulocytosis currently exists. The proposed test of HLA-DQB1 6672G>C has high specificity, but low sensitivity fails to reduce the agranulocytosis risk in the LR group sufficiently that monitoring could be reduced or ceased. Candidate gene studies have failed to identify a strong, replicated genetic variant that substantially increases risk of clozapine induced agranulocytosis.^29, 30^ The first genome-wide association study of clozapine-induced agranulocytosis detected significant associations at two HLA amino acids;^33^ at least one further study is in progress,^53^ and combined analysis of such studies may identify associated genetic variants that can be rapidly translated to clinical practice.

## Acknowledgements

This study was funded by an industrial CASE studentship to Moira Verbelen from the Medical Research Council with Eli Lilly and Company Ltd, by the European Community's Seventh Framework Programme (FP7/2007-2013) under grant agreement n° 279227 (CRESTAR project, http://www.crestar-project.eu/), and under the Marie Curie Industry-Academia Partnership and Pathways, grant agreement n° 286213 (PsychDPC, http://www.psych-dpc.eu/). This study was part-funded by the National Institute for Health Research (NIHR) Biomedical Research Centre at South London and Maudsley NHS Foundation Trust and King’s College London. The views expressed are those of the authors and not necessarily those of the NHS, the NIHR or the Department of Health.

